# Development and evaluation of a non-invasive brain-spine interface using transcutaneous spinal cord stimulation

**DOI:** 10.1101/2024.09.16.612897

**Authors:** Carolyn Atkinson, Lorenzo Lombardi, Meredith Lang, Rodolfo Keesey, Rachel Hawthorn, Zachary Seitz, Eric C. Leuthardt, Peter Brunner, Ismael Seáñez

## Abstract

Motor rehabilitation is a therapeutic process to facilitate functional recovery in people with spinal cord injury (SCI). However, its efficacy is limited to areas with remaining sensorimotor function. Spinal cord stimulation (SCS) creates a temporary prosthetic effect that may allow further rehabilitation-induced recovery in individuals without remaining sensorimotor function, thereby extending the therapeutic reach of motor rehabilitation to individuals with more severe injuries. In this work, we report our first steps in developing a non-invasive brain-spine interface (BSI) based on electroencephalography (EEG) and transcutaneous spinal cord stimulation (tSCS). The objective of this study was to identify EEG-based neural correlates of lower limb movement in the sensorimotor cortex of unimpaired individuals and to quantify the performance of a linear discriminant analysis (LDA) decoder in detecting movement onset from these neural correlates. Our results show that initiation of knee extension was associated with event-related desynchronization in the central-medial cortical regions at frequency bands between 4-44 Hz. Our neural decoder using µ (8-12 Hz), low β (16-20 Hz), and high β (24-28 Hz) frequency bands achieved an average area under the curve (AUC) of 0.83 ± 0.06 s.d. (n = 7) during a cued movement task offline. Generalization to imagery and uncued movement tasks served as positive controls to verify robustness against movement artifacts and cue-related confounds, respectively. With the addition of real-time decoder-modulated tSCS, the neural decoder performed with an average AUC of 0.81 ± 0.05 s.d. (n = 9) on cued movement and 0.68 ± 0.12 s.d. (n = 9) on uncued movement. Our results suggest that the decrease in decoder performance in uncued movement may be due to differences in underlying cortical strategies between conditions. Furthermore, we explore alternative applications of the BSI system by testing neural decoders trained on uncued movement and imagery tasks. By developing a non-invasive BSI, tSCS can be timed to be delivered only during voluntary effort, which may have implications for improving rehabilitation.

## Introduction

Spinal cord injury (SCI) is a life-altering event in which damage to the spinal cord disrupts signals to and from the brain, resulting in lasting motor impairments and paralysis. Rehabilitation strategies such as exercise therapy and gait training can aid in restoring some of the function that was lost with injury, but the effectiveness of these methods tends to plateau within the chronic stage of SCI (> 6 months post-injury) (1–3), and there is presently no cure for paralysis.

Spinal cord stimulation (SCS), applied epidurally and non-invasively during physical therapy, has shown promising effects in improving motor function in the chronic stage of SCI (2,4–6). SCS can provide a temporary prosthetic effect to paralyzed limbs, enabling a wider range of motion in individuals with paralysis, thus aiding in exercise-based programs and promoting functional recovery. Activity-dependent reorganization of spinal circuits enabled by SCS may be a primary contribution towards the restoration of function (2,4,6), where SCS is believed to amplify residual descending voluntary motor input, increasing cortical excitability (6,7) and supporting the excitatory drive to the motoneurons (2,7). Therefore, linking the delivery of stimulation with descending voluntary motor input may result in improved recovery outcomes through rehabilitation.

Brain-computer interfaces (BCIs) in rehabilitation pair feedback with a desired outcome, allowing the user to modulate their cortical activity based on biofeedback and promoting neuroplasticity in different brain regions (8,9). Indeed, neurostimulation techniques that are timed with movement intention lead to better motor outcomes than continuous neurostimulation regardless of movement intention (7,10–13). Invasive brain-spine interface (BSI) systems that link the delivery of SCS to brain-decoded commands have been shown to improve gait function after SCI in proof-of-concept studies involving rats (10), primates (14), and one human (13). However, the invasive nature and higher cost compared to non-invasive alternatives may prevent invasive technologies from helping millions of people living with paralysis. In this work, we present the development and evaluation of a non-invasive BSI using event-related desynchronization in electroencephalography (EEG) to control the delivery of transcutaneous SCS (tSCS) in real time.

The long-term goal of this study is to use this non-invasive BSI to promote recovery of voluntary lower-limb movements in people with SCI. In this work, we developed a neural decoder to detect right knee extension via EEG in unimpaired control participants during a cued movement task. We tested the generalization of the decoder to imagery and uncued movement control conditions. We demonstrated that the neural decoder could predict movement with above-chance accuracy in all conditions, therefore showing that the decoder makes predictions primarily from cortical activity related to movement intention rather than experimental artifacts. The neural decoder was then tested in real-time in brain-controlled tSCS, and we demonstrated that the system could provide an accurate, above chance, closed-loop control of tSCS timed with movement intention. Lastly, we demonstrated potential applications of this BSI system in future rehabilitation contexts in control experiments and additional analyses testing a decoder trained on uncued movement and imagery. These results may have implications for potential rehabilitation methods pairing the delivery of tSCS with the cortical desynchronization of frequency bands commonly associated with movement (15,16), thereby coupling cortical involvement with rehabilitation.

## Methods

### Participant recruitment

This study has been approved by Washington University in St. Louis’ Institutional Review Board (IRB ID 202105168). Seventeen unimpaired participants (10 male, 7 female, average age 25.8 ± 3.9 years) were recruited for this study. Participants performed one or more of three phases: i) offline decoder validation without tSCS, ii) real-time decoder testing with brain-controlled tSCS, or iii) control conditions. Participant demographics and the phases they completed are summarized in **Supplementary Table 1**.

### Data acquisition and processing

Electroencephalography (EEG) data were recorded at a 500 Hz sampling rate using a wireless 32-channel EEG headset (gNautilus, gTec, Austria) and base station (gNautilus Base Station, gTec, Austria) within the BCI2000 software (17) (**Figure 1a**). The 32 channels were positioned with a 64-channel electrode density over the central-medial areas according to the 10-10 system (18) to record relevant signals over the sensorimotor cortex.

**Figure 1:**
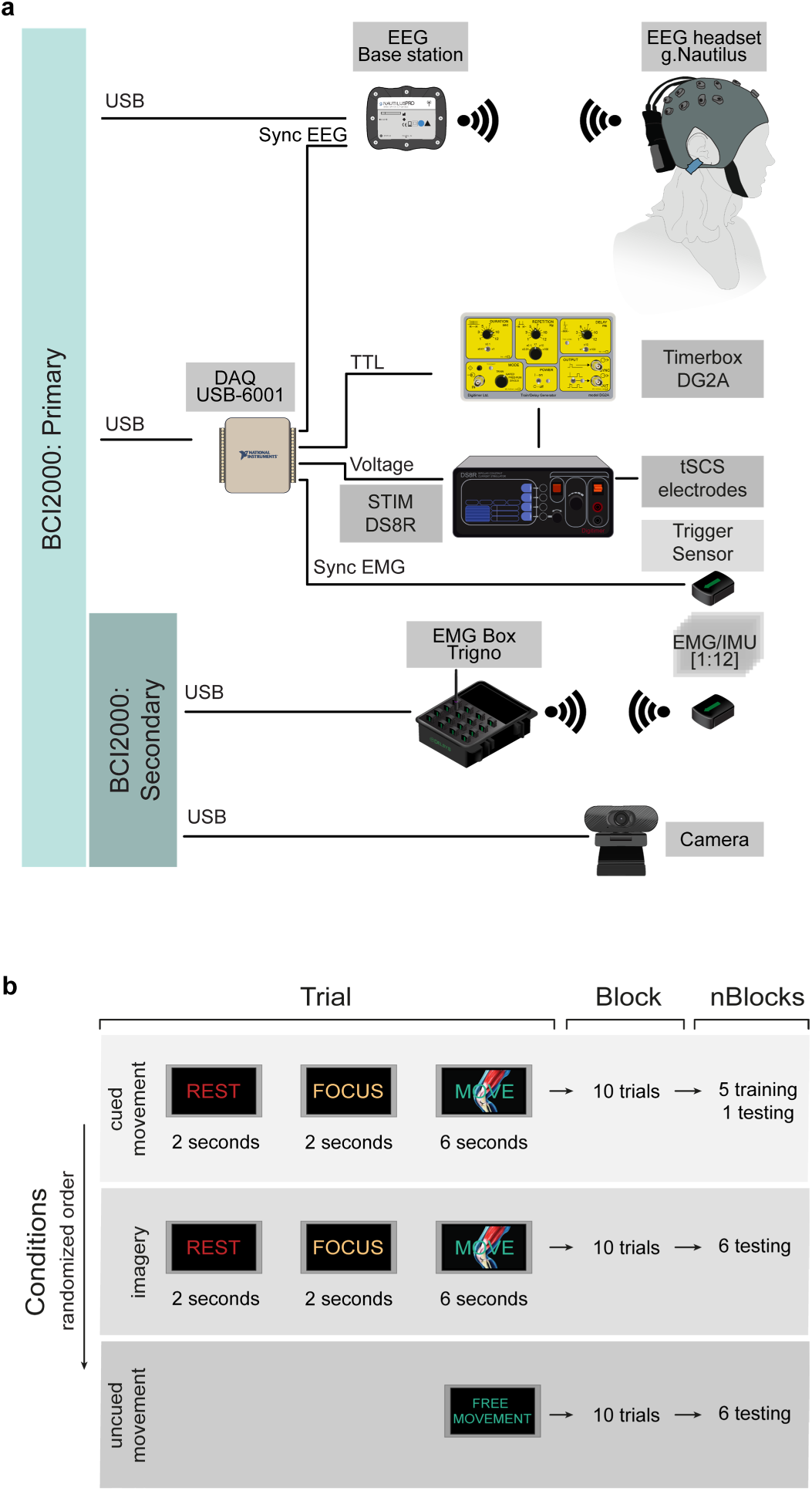
Technological framework and experimental design. **a.** Hardware setup. Technological framework for EEG and EMG signal acquisition and the delivery of tSCS using BCI2000 (17). **b.** Phase I experimental protocol. Phase I consisted of 3 conditions, 6 blocks per condition, and 10 trials (repetitions) per block. Cued movement blocks were used to train the decoder with 5-fold cross-validation, with the sixth block reserved for testing.

Electromyography (EMG) signals were recorded at a 1482 Hz sampling rate using wireless surface electrodes (Trigno® Avanti, Delsys Inc., USA) placed bilaterally over the rectus femoris, vastus lateralis, tibialis anterior, medial gastrocnemius, and soleus muscles according to the SENIAM convention (19). Before placing the EMG sensors, skin was prepped using abrasive gel (NuPrep®, Weaver and Co. USA) with a cotton swab and then wiped off with alcohol pads. A secondary BCI2000 instance logged the EMG/IMU signals along with a participant video at 30 fps (C270 HD Webcam, Logitech, Switzerland). A NI-DAQ (DAQ USB-6001, National Instruments, USA) digital input/output board generated a square sync pulse that was fed to the EEG headset base station and a dedicated analog sensor (Trigno Avanti Analog, Delsys, USA) to ensure consistent synchronization between data streams. The DAQ controlled stimulation amplitude and a digital output set to trigger a biphasic constant current stimulator (DS8R, Digitimer, UK) through a digital pulse train generator (DG2A, Digitimer, UK) (**Figure 1a**).

### Experimental setup

Participants sat approximately 1 m in front of a 1.5 x 2.3 m projector screen and were asked to perform a right knee extension in response to a visual-auditory cue delivered according to one of three experimental phases. The objectives of the first phase (*Phase I: Neural correlates of knee extension*) were to i) identify the neural correlates of lower-limb movements in unimpaired participants, ii) develop a linear discriminant analysis (LDA)-based decoder of knee extension, and iii) assess its ability to predict knee extension offline. The objectives of the second phase (*Phase II: Brain-controlled stimulation*) were to evaluate the ability of the LDA-based decoder to i) predict leg extension online and ii) control the delivery of tSCS based on brain-decoded commands. The objectives of the third phase (*Phase III: Controls and alternative strategies)* were to i) perform control conditions to ensure the neural decoder was truly learning from cortical activity related to leg extension, ii) investigate the neural mechanisms of movement onset predictions, and iii) demonstrate potential alternative applications of a BSI system in SCI rehabilitation contexts.

### Phase I: Neural correlates of knee extension

Participants were instructed to extend their right knee following a visual-auditory cue (**Figure 1b**, cued movement condition), imagine extending their right knee without moving while following the same cue (**Figure 1b**, imagery condition), or extend their right knee several times without cues using their preferred timing (**Figure 1b**, uncued movement condition). The cue consisted of a sequence of “*Rest*”, “*Focus*”, and “*Move*” periods during which these words were displayed on the screen. During the “*Move*” period, a video of an anthropomorphic knee extension was displayed. The imagery condition was used as a control for movement-related artifacts and to gain insights into potential applications in cases of complete SCI, in which no residual movement could be used to label training data. The uncued movement condition was used to control for cue-related artifacts and to determine the ability of the BSI to function independently without cues. To reduce variability across conditions, participants were instructed to perform the same slow movements during cued and uncued movement conditions. Conditions were performed in block-randomized order, with each block consisting of ten repetitions of leg extension, and six blocks performed per condition (**Figure 1b**).

### Data processing, feature extraction, and LDA decoder training

Raw EEG signals were bandpass filtered between 4-40 Hz to extract frequency bands of sensorimotor rhythms (15,20,21), common average referenced to remove movement artifacts (20–22), and further decomposed in frequency in 4 Hz bins using a 4th-order Butterworth bandpass filter bank (**Figure 2c**). Power spectral density (PSD) was calculated by squaring the amplitudes, low-pass filtering at 2 Hz with a 4th-order Butterworth filter, and dividing by the bandwidth of the bin (21). The filtered signal was then down-sampled to 10 Hz to improve decoder stability.

**Figure 2:**
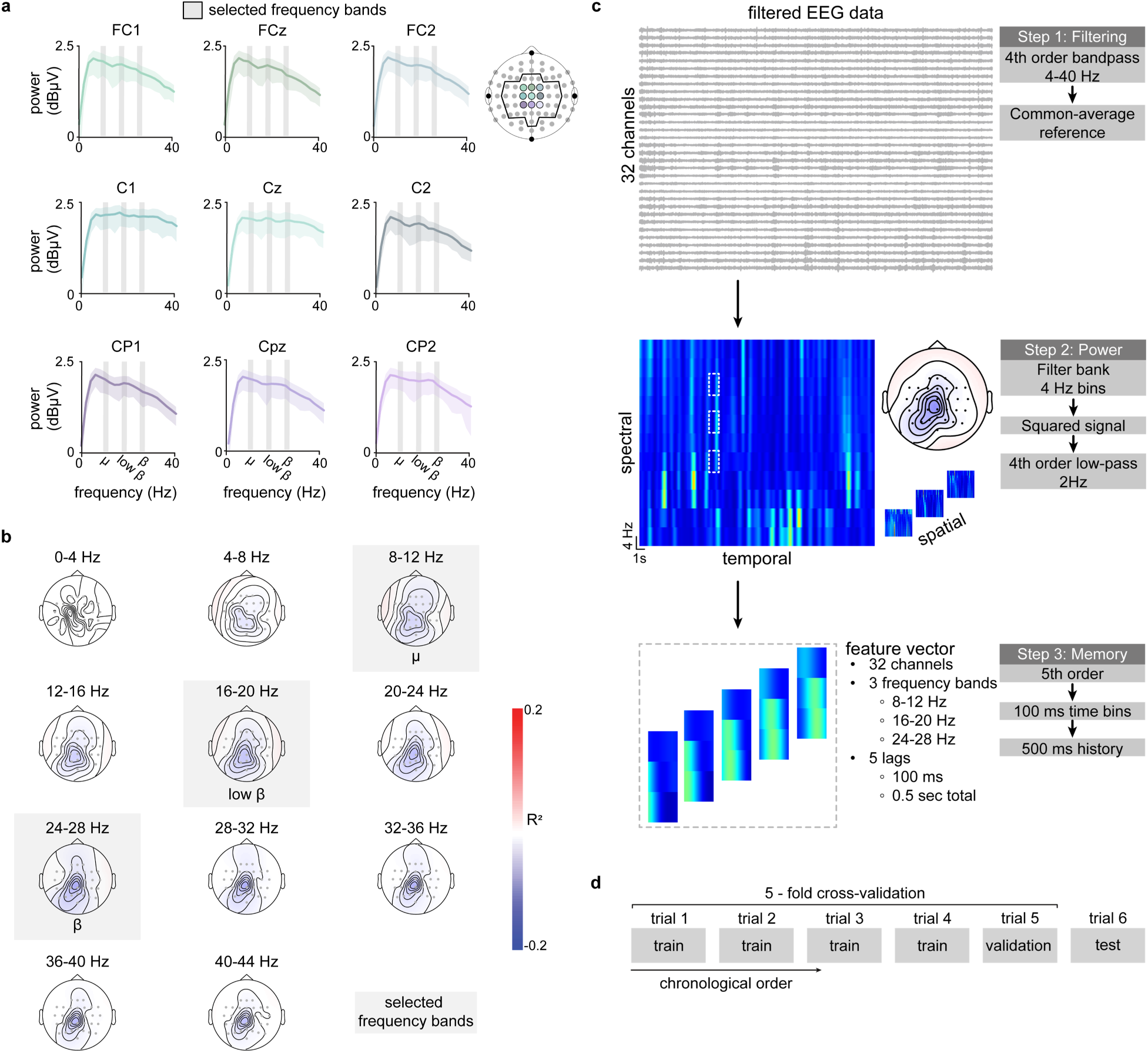
Analysis of predictive frequency bands informed selection of feature space used in the LDA decoder. **a.** Power spectral data recorded via sensorimotor channels during right knee extension for a representative participant. Analysis of the power spectral data during movement of pilot participants informed the selection of frequency bands within the feature space. **b.** R^2^ scalp topographies for a representative participant. R^2^ was computed between the true movement label, and the power spectrogram was computed for each channel. Pilot data revealed sensorimotor desynchronization in several frequency bands, including µ (8-12 Hz), low β (16-20 Hz), and high β (24-28 Hz). Non-neighboring frequency bands below 30 Hz were selected to prevent overlap in information fed into the decoder and to avoid the stimulation artifact at 30 Hz with the future addition of real-time tSCS. **c.** EEG data processing pipeline. EEG data was bandpass 4-40 Hz filtered and common average referenced. Power was extracted in 4 Hz bins by band- passing, squaring, and low-pass filtering the common average referenced data. 480 features were extracted corresponding to 3 frequency bands (µ, low β, and high β), 5 lags, and 32 channels. Lags were incorporated so movement onset predictions can take data from the past 0.5 seconds into account. **d.** Five-fold cross-validation decoder training. The decoder was trained with a 5-fold cross-validation strategy in which four blocks were used as training blocks. Once the hyperparameters were optimized to minimize validation error, the model was retrained on all five blocks and tested on the sixth, unseen block.

Time-frequency domain features were extracted from the PSDs and used to train the LDA classifier using the µ (8-12 Hz), low β (16-20 Hz), and high β (24-28 Hz) frequency bands. These frequency bands were selected based on our preliminary analysis, which showed that power fluctuations in the 4-44 Hz frequency band explain a large fraction of movement-related variance (**Figure 2a, b**) and previous reports indicating that these frequency bands contain important information related to sensorimotor function (6,12,15,16,21). We avoided the 30 Hz band to prevent contamination by stimulation artifacts from tSCS in Phase II.

A two-class LDA decoder was trained based on 480 extracted features corresponding to spectral (3 frequency bands), temporal (five previous 100 ms windows), and spatial (32 channels) information (**Figure 2c**) to distinguish between movement and no movement. This approach maintained an approximate 1:10 ratio between the number of features and the size of the expected training dataset (23), reducing the risk of overfitting. Ground truth movement data was determined from the cue to compare the cue vs. imagery conditions and from the IMU to compare the cue vs. uncued movement conditions. Movement onset from the IMUs was determined from an empirically tuned threshold crossing of angular velocity. The training data was randomly up- sampled to ensure class balance, and the seed was fixed for reproducibility. The neural decoder was trained on the first 5 blocks with a 5-fold cross-validation strategy to optimize the hyperparameters (**Figure 2d**). A covariance regularization term and a zero-rounding threshold were selected as the hyperparameters that minimized classification error within the 5-fold cross-validation. The decoder was then retrained using the entire training set and tested on the 6^th^ (last, unseen) block of each condition to minimize the risk of temporal leakage and to account for degraded performance over a session related to changes in EEG signal quality. All performance metrics reported refer to the test set.

To evaluate how the spatial and spectral data changed between rest and movement, topographical R^2^ was calculated for pilot participants during the cued movement condition. Signed R^2^ was computed as the coefficient of determination between the true movement label and the power spectrograms of each channel and thus represented how much of the overall variance in power for each channel can be attributed to movement (24).

### Phase II: Brain-controlled stimulation

Participants first performed six cued movement blocks with artificially controlled stimulation to train, validate, and test the neural decoder. These blocks were followed by six additional cued movement blocks to evaluate the brain-controlled stimulation in real time. The decoder was then re-trained on all cued movement blocks and tested for generalization of brain-controlled stimulation during uncued movement.

Transcutaneous spinal cord stimulation (tSCS) was applied with a 3.2 cm diameter cathode located to the right of spinal segment T10 and a 7.5 x 10 cm return electrode on the right side of the navel to target the right leg proximal muscles as previously described in our work (25) (**Supplementary Figure 1**). During the first six blocks of decoder training, stimulation was delivered at 30 Hz. The amplitude was ramped up from 0 to 10 mA during the first 7 seconds of each block and alternated between 10 mA during the rest periods and 15 mA during the movement periods. Although 15 mA was generally below the motor threshold, we chose this stimulation amplitude to help with participant retention, as continuous 30 Hz stimulation at high intensities can be difficult to tolerate for unimpaired individuals with intact sensory function. The stimulation protocol alternating between 10-15 mA was developed i) to improve participant comfort by preventing large jumps in stimulation amplitude between the rest and movement periods and ii) to ensure that neurophysiological effects and possible stimulation artifacts were accounted for so that the decoder could learn to predict movement intention in the presence of tSCS. The neural decoder was trained using a 5-fold cross-validation strategy, and two cued movement runs were used to empirically tune the probability threshold while controlling the stimulation from brain-decoded commands in real time. This manual tuning aimed to align stimulation with movement onset while balancing true/false positives and incorporating the participants’ verbal feedback on the perceived alignment of stimulation.

The modified decoder was used to trigger stimulation onset in real time during the next six blocks of cued movement. Stimulation was triggered (increased from 10 mA to 15 mA) when the calculated probability rose above the probability threshold and was left on (15 mA) for a fixed duration of 6 seconds to match the expected movement cue duration, as done in previous work on brain-controlled spinal cord stimulation in non-human primates (14). In addition, stimulation was re-set to baseline (10 mA) at the end of each movement cue with a 1-second refractory period to prevent the 6-second stimulation from continuing into the rest period. The decoder was then re-trained using an 11-fold cross-validation and was tested for generalization to two blocks of uncued movement.

### Phase III: Controls and alternative strategies

Action observation has been shown to activate motor regions related to the observed movement (26). To account for possible confounds introduced by using an anthropomorphic cue, we evaluated decoder generalization performance when two participants trained with and without the anthropomorphic cue (**Supplementary Table 1**). Participants performed six blocks of cued movement using a rectangle with changing height as the movement cue (bar cue), followed by six blocks of cued movement using the anthropomorphic knee flexion graphic used in Phases I and II (knee cue). Decoders were tested for generalization to the uncued movement condition. These conditions were tested without tSCS. These conditions were not randomized; the bar cue was presented first to prevent participants from recalling the visual image of the anthropomorphic cue.

To account for differences in generalization performance due to possible alternate neural strategies between cued and uncued movement, we evaluated the real-time performance of a neural decoder trained and tested on uncued movement in a group of four participants (**Supplementary Table 1**). Participants performed six uncued movement runs with artificially controlled stimulation for decoder training. During the training blocks, stimulation amplitude alternated between rest (10 mA) and movement (15 mA) based on a threshold crossing on the shin’s IMU pitch angle. The decoder was then trained using 5-fold cross-validation as previously described and tested in real-time on an additional six uncued movement runs with brain-controlled stimulation.

### EEG noise and participant exclusion

In all experiment phases, EEG channel noise was evaluated offline and used as a rejection criterion for participant data. The median of each channel’s rectified signal was compared to all channels’ interquartile range. Channels were considered noisy when their median was outside the interquartile range by a factor of three (i.e. < Q1 – 3*IQR or > Q3 + 3*IQR). As noisy channels greatly impacted common average referencing, blocks with any single noisy channel were removed. Two participants excluded from the analysis had noisy channels across all blocks throughout the experiment (**Supplementary Table 1**).

### Analysis and performance metrics

Decoder performance was calculated for each participant by computing the receiver operating characteristic (ROC) curve and finding the area under the curve (AUC). The AUC was chosen as the performance metric since it is agnostic of the probability threshold and robust to class imbalance. All conditions were compared against a chance AUC of 0.5.

To evaluate how the spatial and spectral data changed between the imagery and uncued movement conditions compared to the cued movement condition, topographical R^2^ was calculated for all participants across conditions. Each channel’s R^2^ during the imagery and uncued movement condition was then subtracted from the respective channel’s R^2^ in the cued movement condition. The differences in R^2^ were plotted topographically for visualization. In addition, we performed a principal component analysis (PCA) of the 32-channel R^2^ data. Principal components were calculated in the same space for all conditions and participants but separately for each frequency band. We quantified the Euclidian distance between each participant’s neural state (27) during the cued movement vs. imagery or uncued movement conditions along the first three principal components.

We argue that perception of neural decoder performance would vary according to a participant’s tolerance for time discrepancies between their movement and stimulation onset. To understand the relationship between performance and tolerance, we quantified decoder performance as a function of a tolerance window around true movement onset. Predicted onsets were determined based on a positive crossing of a probability threshold. The threshold was set to the empirically tuned thresholds used online in Phase II and to 0.73 in Phase I, which was the average of the tuned thresholds across Phase II. True positives and negatives were determined from predicted onsets within the tolerance window, and these were calculated for tolerance windows between 0 and 3 seconds.

### Statistics

The 95% confidence intervals on the AUC for each participant were computed by calculating performance on bootstrapped (sampling with replacement) test data over 500 iterations. Confidence intervals for all participants and conditions are reported in **Supplementary Table 3**. The AUCs used in group analysis were not part of the bootstrapped data but were based on the performance obtained from the original testing datasets. AUCs were statistically compared across experimental conditions using a non-parametric paired samples Wilcoxon signed rank test and compared to a chance AUC of 0.5 using a one-sample Wilcoxon signed rank test.

In Phase III, we used bootstrapping to compare decoder performance between groups and conditions with a different number of participants (7,28). The populations were sampled with replacement to obtain k = 10,000 fictive populations to simulate a distribution for each condition. We then tested the null hypothesis that there was no significant difference between the means for both distributions. A p-value was calculated by estimating the overlap of the residuals with a Bonferroni correction for multiple comparisons. On the dataset with a small number of participants (n = 4), the simulated bootstrapped distribution was used in the comparison to chance.

R^2^ differences between conditions were statistically compared to zero using a one-sample signed rank test. Due to the high number of electrodes (32 channels) and low number of participants (7 participants), this analysis was not corrected for multiple comparisons. Correcting for multiple comparisons would reduce the power to detect meaningful effects, increasing the risk of Type II errors; therefore, results are presented without correction while acknowledging this limitation.

## Results

### EEG-based neural decoder can reliably predict movement during cued knee extension offline

In the first phase of this study, we asked whether the LDA decoder trained on the EEG data collected during the cued knee extension task could reliably predict movement onset offline better than chance. The extracted EEG features synchronized with EMG, knee angle, and the calculated movement probability for a representative participant during a single testing block are shown in **Figure 3a**. The movement probability generally increased between the focus and movement phases (**Figure 3d**). Group analysis revealed an average AUC of 0.83 ± 0.06 s.d. across seven participants, which was significantly higher than chance (**Figure 3g**, Hedge’s g = 4.29, p = 0.02*, n = 7). Moreover, decoder performance was evaluated by examining the accuracy of predictions at every timepoint with a probability threshold of 0.73 and calculating a confusion matrix (**Figure 3j**). This resulted in an average true positive rate (TPR) of 92% and a true negative rate (TNR) of 54%, implying that, on average, the probability predictions were correct for 73% of timepoints. These results indicate that a decoder trained and tested on cued movement can reliably predict movement onset from the extracted feature space.

**Figure 3:**
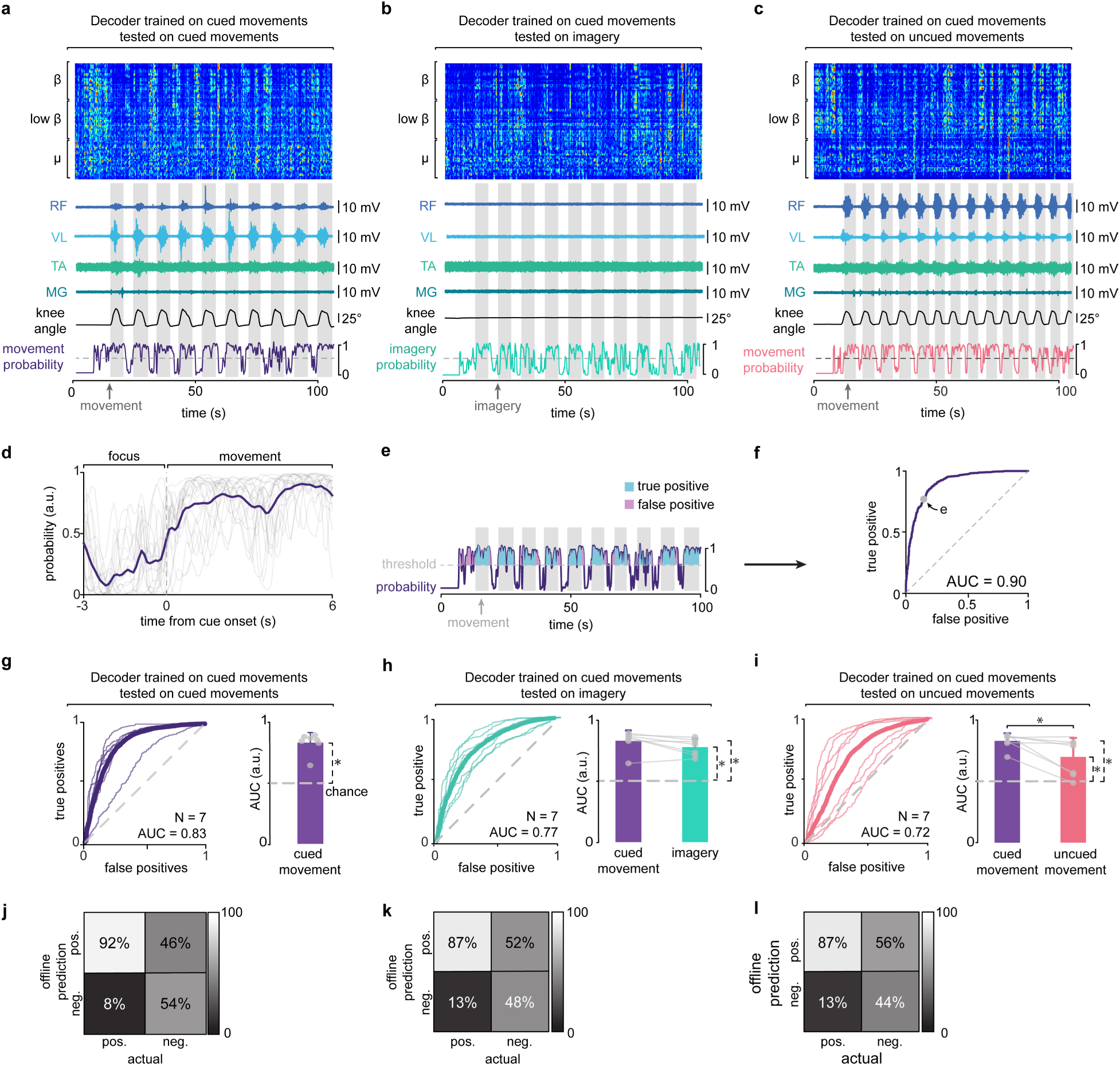
Offline decoder performs above chance for cued, imagery, and uncued movement. **a.** EEG spectrograms aligned with EMG and movement kinematics during cued movement. A single participant’s EEG power features during offline testing on cued movements decomposed by frequency band aligned in time with EMG signals from the vastus lateralis (VL), rectus femoris (RF), tibialis anterior (TA), and medial gastrocnemius (MG), knee angle, and probability calculated offline. Desynchronization was observed in the EEG spectrograms across frequencies at the onset of movement. **b.** EEG spectrograms aligned with EMG and movement kinematics during imagery. **c**. EEG spectrograms aligned with EMG and movement kinematics during uncued movement. **d.** Probability for single trials (thin lines) and averaged across trials (thick line) during focus and movement periods. Movement probability increased after movement onset. **e.** Illustration of true positive and false positive rate calculation from probability. True positive and false positive rates are calculated at a sweep of probability thresholds between 0 and 1 and used to construct a receiver operating characteristic (ROC) curve. **f.** ROC curve for a single participant. The ROC curve was calculated for a single participant by varying the threshold on the probability and comparing to true movement. The probability threshold used in **e** is shown on the ROC curve. The area under the ROC curve (AUC) was used to quantify all decoder performances. **g.** ROC curve for each participant (thin lines) and averaged across participants (thick line) when testing on cued movement and average area under ROC curve (AUC) compared to chance. **h.** Same as **g** but for a decoder tested on imagery. Paired comparison between cued and imagery and comparison of each to chance. **i.** Same as **g** but for a decoder tested on uncued movements. Paired comparison between cued and uncued movement and comparison of each to chance. **j.** Confusion matrix at a fixed probability threshold when tested on cued movement. **k.** Same as **j** but for a decoder tested on imagery. **l.** Same as **j** but for a decoder tested on uncued movement. Bars in **g-i** represent mean ± s.d., with each circle representing the testing AUC for each participant. The asterisks on the right of each bar represent the results of the one-sample Wilcoxon signed rank test for each decoder’s AUC against chance; the asterisks between bars represent the paired samples Wilcoxon signed test between two decoder’s average AUC. * p < 0.05; ** p < 0.01; *** p < 0.001. Abbreviations: rectus femoris (RF), vastus lateralis (VL), tibialis anterior (TA), medial gastrocnemius (MG).

### Generalization to imagery and uncued movement suggest robustness against movement and cue-related artifacts

The spectral EEG data during the imagery and uncued movement conditions showed movement-related desynchronization, and offline-calculated probability was generally aligned with the movement phases (**Figure 3b, c**). A group analysis on the performance of the cued movement-based decoder tested on imagery blocks revealed an average AUC of 0.77 ± 0.07 s.d., which was not significantly different from the performance of the same decoder tested on cued movement (**Figure 3h**, Hedge’s g = 0.73, p = 0.08, n =7). Additionally, this decoder’s performance was significantly above chance (Hedge’s g = 3.68, p = 0.02*). Confusion matrices averaged across participants revealed a TPR of 87% and TNR of 48%, indicating that on average, predictions were correct for 68% of timepoints when tested on imagery.

The group analysis for the cued movement decoder tested on the uncued movement blocks revealed an average AUC of 0.72 ± 0.15 s.d., which was significantly lower than cued movement performance (**Figure 3i**, Hedge’s g = 1.01, p = 0.03*, n = 7). However, this performance was still significantly higher than chance (Hedge’s g = 1.48, p = 0.03*). Confusion matrices across participants resulted in a TPR of 87% and TNR of 44%; thus, on average, 67% of time points were correctly predicted when tested on uncued movement.

Although performance slightly decreased when the decoder trained on the cued movement condition was tested for generalization to the imagery and uncued movement conditions, this performance was higher than chance. This suggests that although movement- and cue-related artifacts may have a small impact on decoder performance, the successful decoding performance of the cued movement-based decoder cannot solely be attributed to these artifacts. Moreover, these differences in performance could be attributed to intrinsic differences in neural strategies between tasks. Therefore, we next sought to investigate differences in desynchronization patterns across tasks.

### Desynchronization patterns during cued movement have focused differences with imagery but global differences with uncued movement

To investigate differences in neural strategies between tasks, we compared topographical R^2^ maps across conditions. Topographical R^2^ maps for two representative participants are shown in **Figure 4a**. These participants were selected as representative examples of participants with good (S005) and poor (S001) generalization to the uncued movement condition, as denoted by the AUC. Note that the topographical R^2^ pattern in S005 is quite similar across conditions, and this participant had an AUC of 0.80 and 0.82 for the imagery and uncued movement conditions, respectively. In contrast, while the R^2^ pattern between imagery and cued movement was similar in S001, the pattern looks markedly different during the uncued movement condition. This participant had an AUC of 0.86 for the imagery condition and an AUC of 0.58 for uncued movement.

**Figure 4:**
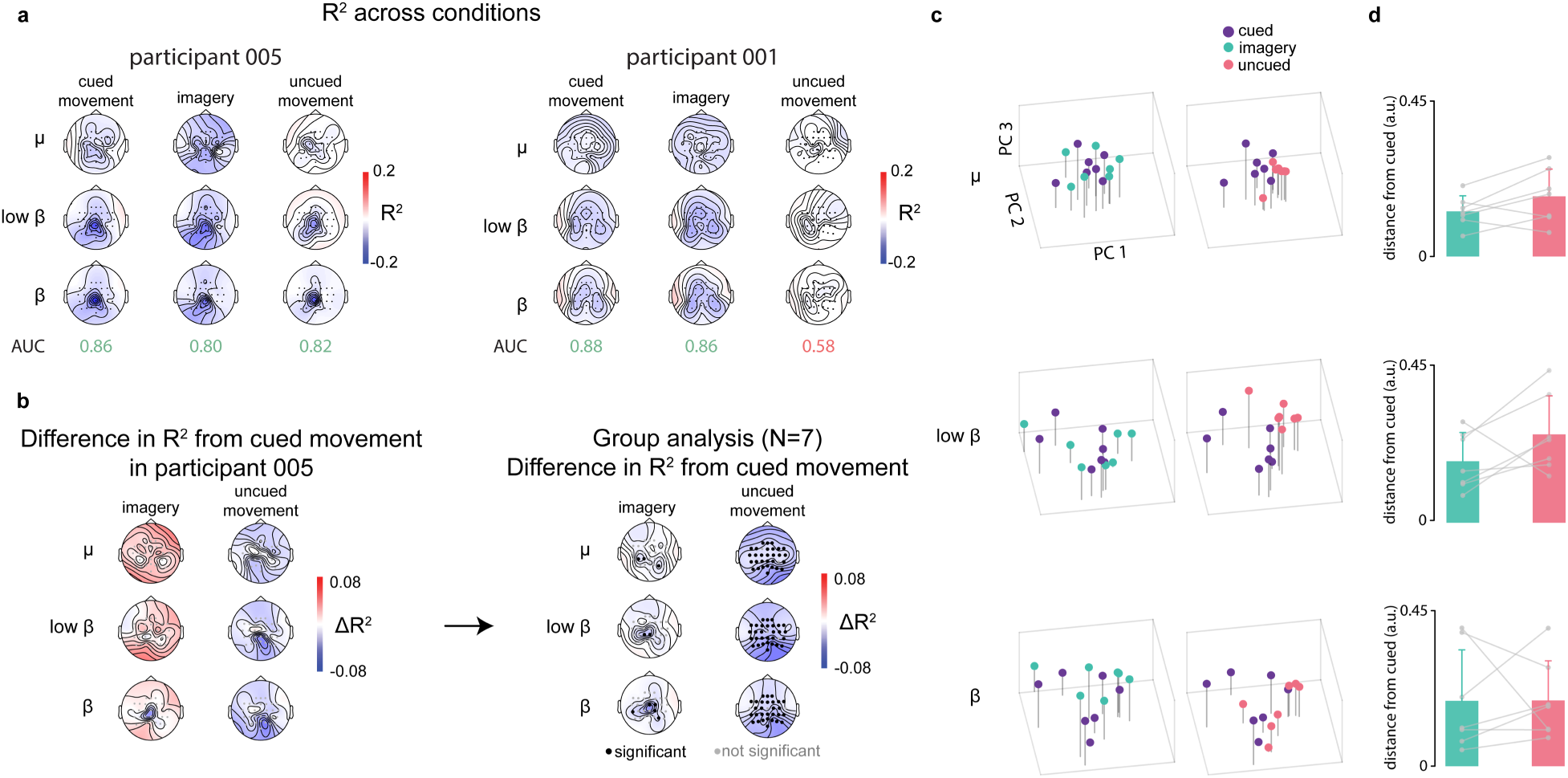
Analysis of R^2^ suggests spectral and spatial differences in EEG activity between conditions. **a.** R^2^ scalp topography plots during movement for two participants showing good (left, participant S005) and poor (right, participant S001) generalization across conditions. Consistencies in spatial and spectral activity across conditions resulted in better decoder generalization evidenced by higher AUCs. **b.** Difference in R^2^ during movement between conditions for one participant (left) and averaged across participants (right). Group analysis of R^2^ differences between conditions revealed focused differences between cued movement and imagery, and widespread differences between cued and uncued movement. Significant channels were not corrected for multiple comparisons. **c.** PCA projections of R^2^ scalp topographies for all conditions and frequency bands. R^2^ data across electrodes was projected onto the first three PCs. Each point represents one participant, and data is color-coded by condition. PCs across conditions (left and right columns) are the same. Conditions shown separately for visualization purposes. **d.** Average Euclidean distance of imagery and uncued movement to cued movement in PC space. Distance between uncued and cued movement is slightly larger than the distance between imagery and cued movement, but this effect is not significant.

The topographical maps for differences in R^2^ patterns between cued movement and the imagery and uncued movement conditions for participant S005 and averaged across participants are shown in **Figure 4b**. The group analysis revealed that while differences in desynchronization between cued movement and imagery were focused on the central median channels, differences between cued and uncued movement were widespread across all channels.

Unlike classification algorithms such as LDA, principal components in PCA are not selected to separate different classes but to explain the most variance. Nevertheless, the neural states for the different conditions were somewhat isolated in PC space (**Figure 4c**). The distance in PC space between the uncued movement and cued movement conditions was generally higher than that between the imagery and cued movement conditions (**Figure 4d**). However, this difference was not statistically significant.

### Real-time brain-controlled tSCS in unimpaired individuals

We tested the real-time performance of delivering tSCS timed with movement intention predicted from the extracted EEG features (**Figure 5a**). Stimulation was set to a baseline of 10 mA and increased to 15 mA during movement, modulated by the task cue in the training set (**Figure 5b**). In real-time testing, desynchronization of the sensorimotor channels was synchronized with movement onset shown by leg muscle EMGs and movement kinematics (**Figure 5c**) and was used by the decoder to control the onset of stimulation.

**Figure 5:**
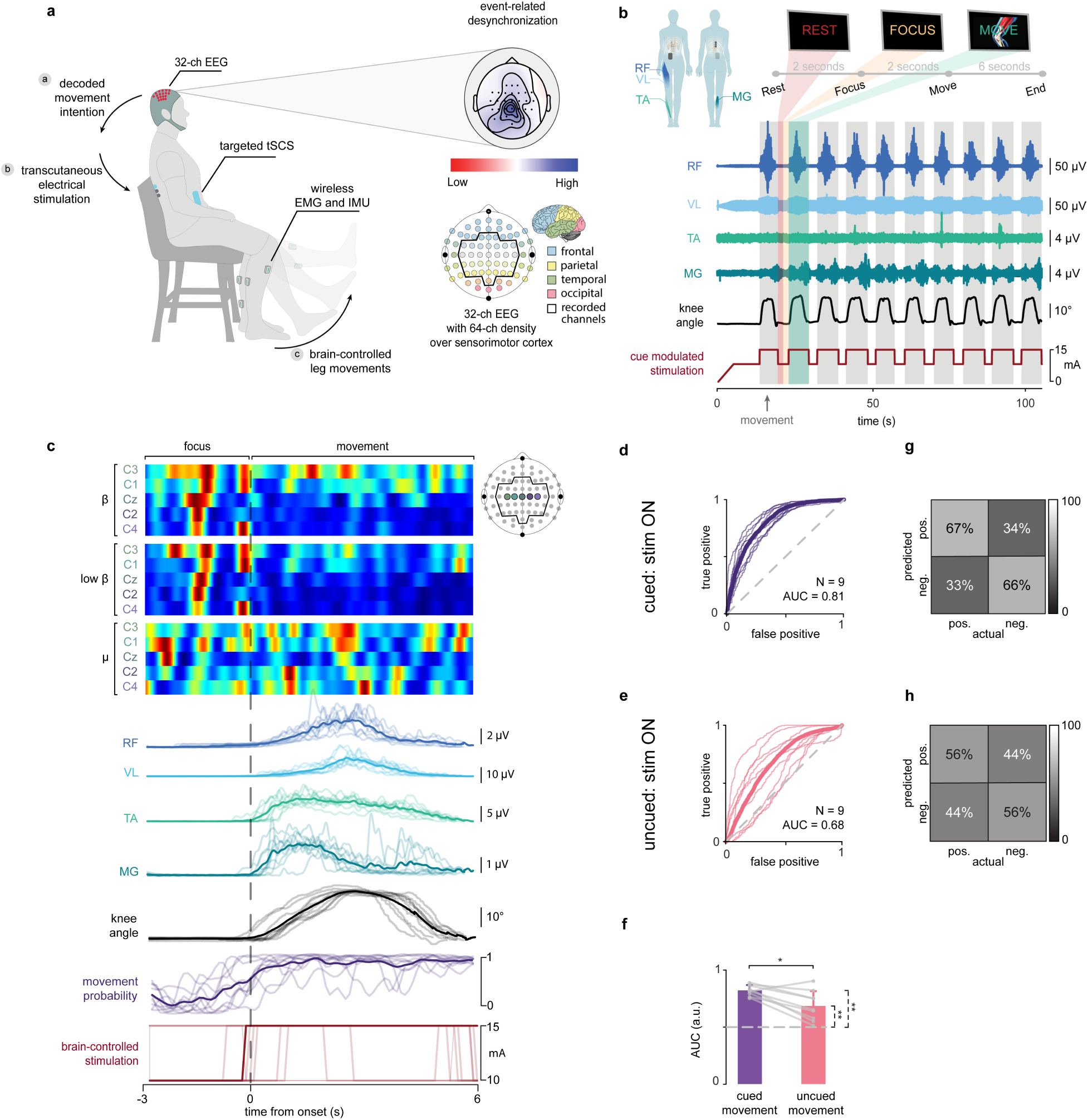
Real-time, closed-loop control of tSCS using predictions of movement intention in a non-invasive BSI. **a.** Technological framework for non-invasive BSI. Desynchronization in the sensorimotor cortex was identified using a 32-channel EEG system in real-time as participants extended their right knee. The predicted movement intention was used to trigger the delivery of tSCS at a higher amplitude. **b**. Illustration of a cued movement block used in the training set. Stimulation was ramped up to a baseline of 10 mA and was increased to 15 mA during the movement phases using the task’s movement cue. **c.** EEG spectrograms of selected sensorimotor channels, kinematics, movement probability, and real-time closed-loop stimulation for a single block for a representative participant (thin lines) and averaged across trials (thick line). Note that there is event-related desynchronization before and after movement onset. Stimulation onset was generally timed with movement onset as shown by the stimulation averaged across trials. **d.** ROC curve for individual participants (thin lines) and averaged across participants (thick line) when testing on cued movement with stimulation. **e.** Same as **d** but for a decoder tested on uncued movement with stimulation. **f.** Paired comparison of average AUC across participants for cued and uncued movement and comparison to chance. **g.** Confusion matrix calculated according to the stimulation administered during the trial and averaged across participants. **h.** Same as **g** but for a decoder tested on uncued movement. Bars in **f** represent mean ± s.d., with each circle representing the average AUC for one participant. The asterisks on the right of each bar represent the results of the paired samples Wilcoxon signed test for each decoder’s AUC against chance; the asterisks between bars represent one-sample Wilcoxon signed test between two decoders’ average AUCs. * p < 0.05; ** p < 0.01; *** p < 0.001. Abbreviations: rectus femoris (RF), vastus lateralis (VL), tibialis anterior (TA), medial gastrocnemius (MG). EEG map and brain region division modified from (29).

Group analysis revealed an average AUC of 0.81 ± 0.05 s.d. when the decoder was tested on cued movement with stimulation (**Figure 5d**) and an average AUC of 0.68 ± 0.12 when tested on uncued movement with stimulation (**Figure 5e**). Consistent with Phase I results, the performance on brain-controlled tSCS during uncued movements was significantly lower than the performance on cued movements (Hedge’s g = 1.28, p = 0.01**, n = 9). However, the performance on both conditions was significantly above chance (**Figure 5f**; Hedge’s g = 6.51, p = 0.004** tested on cued movements, and Hedge’s g = 1.33, p = 0.004** tested on uncued movements, n = 9).

We examined the stimulation paradigm’s accuracy at every timepoint by constructing a confusion matrix based on the real-time administered stimulation compared to true movement. This resulted in an average TPR of 67% and TNR of 66% for cued movement across participants (**Figure 5g**) and a TPR of 56% and TNR of 56% for uncued movement, implying that on average, stimulation was correctly on or off for 67% and 56% of the cued and uncued blocks, respectively (**Figure 5h**). Together, our results demonstrate the feasibility of an LDA-based neural decoder to predict movement intention and control tSCS in real-time.

### Training on uncued movement does not significantly improve decoder performance in uncued movement

Due to the marked changes in neural strategy between cued and uncued movement, we asked whether training a decoder in a condition that resembles the desired application would improve performance. Group analysis (n = 4) for the decoder trained on uncued movement revealed an average AUC of 0.74 ± 0.09 s.d (**Figure 6a**). While the average AUC increased, the performance was not significantly different than a decoder trained on cued movement (**Figure 6b**; Cohen’s d = 1.56, p = 0.12). A confusion matrix was calculated from the real-time stimulation and averaged across participants. The TPR was 50% and TNR was 55%, implying that stimulation was correctly on or off for 53% of the block (**Figure 6c**).

**Figure 6.**
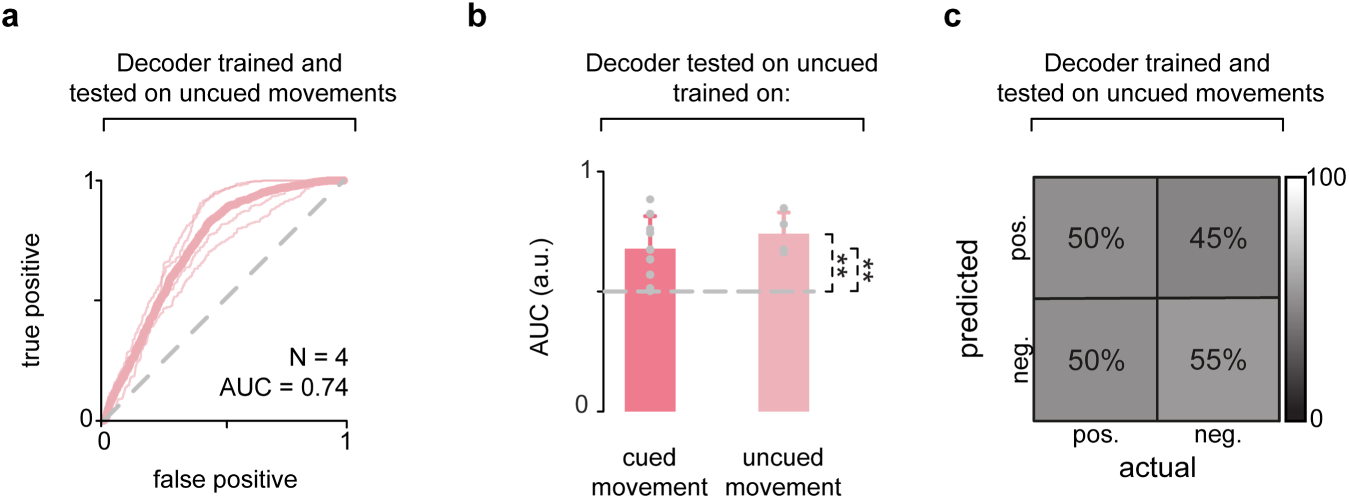
Cued and uncued movement-trained decoders’ performance on uncued task. **a.** ROC curve for all participants (thin lines) and averaged across participants (thick line) when the decoder was trained and tested on uncued movement. **b.** Average AUCs for a decoder tested on uncued movement trained on cued and uncued movement. The performance of the decoder trained on uncued movement (n = 4) was generally higher than the performance of the decoder trained on cued (n = 9), but not significantly different when compared with a bootstrapping analysis. **c.** Confusion matrix calculated from the real-time stimulation and averaged across participants. The mean of TPR and TNR was 53%. Bars in **b** represent mean ± s.d., with each circle representing the average AUC for one participant. The asterisks on the right of each bar represent the results of a one-sample Wilcoxon signed rank test for each decoder’s AUC against chance. * p < 0.05; ** p < 0.01; *** p < 0.001.

### Training on imagery can be used for generalization to movement conditions

People with complete SCI may not be able to generate a movement that could be measured by the EMG/IMUs to train the decoder. To overcome this limitation, the decoder used in the real-time control of tSCS would need to be trained on data obtained during motor imagery. The offline performance of the decoder trained on imagery and tested on cued movement is shown in **Figure 7a**. Group analysis (n = 7) revealed an average AUC of 0.79 ± 0.09 s.d. This performance was not significantly different from the decoder’s performance when trained on cued movement (Hedge’s g = 0.50, p = 0.29), and both were significantly above chance (**Figure 7b**; Hedge’s g = 3.21, p = 0.02* for imagery-based decoder and Hedge’s g = 4.23, p = 0.02* for cued movement-based decoder, n = 7). The confusion matrix averaged across participants resulted in a TPR of 74% and TNR of 43%, implying that when applied, stimulation would be correctly on or off for 59% of the block on average (**Figure 7c**). Therefore, while a decoder can generalize to imagery when trained on cued movement (Phase I), it can also generalize to cued movement when trained on imagery. These results are in agreement with previous reports that imagery-based decoders can be used to control BCI devices in real-time (30–35) and suggest that our system could potentially be applied in cases of complete SCI.

**Figure 7.**
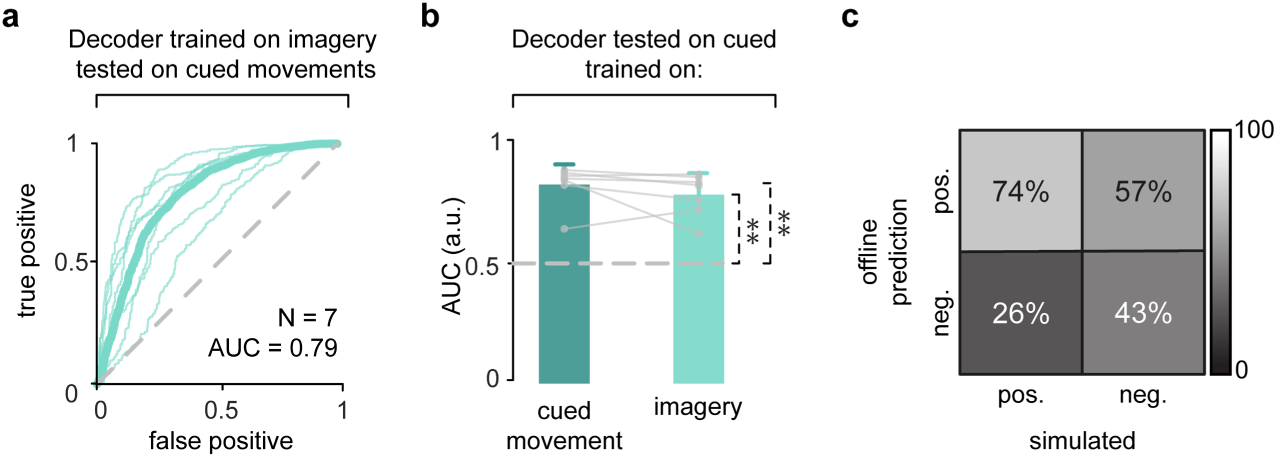
Imagery-trained decoder can generalize to cued movements. **a.** ROC curve for all participants (thin line) and averaged across participants (thick line) when the decoder was trained on imagery and tested on cued movement. **b**. Paired comparison of average AUC across participants for a decoder tested on cued movement, trained on cued movement and imagery. AUC between decoders trained on cued and imagery were not significantly different and were both significantly higher than chance. **c.** Confusion matrix calculated from simulated stimulation according to the implemented stimulation paradigm and averaged across participants for a decoder trained on imagery and tested on cued movement. The mean of TPR and TNR was 59%. Bars in **b** represent mean ± s.d., with each circle representing the average AUC for one participant. The asterisks on the right of each bar represent the results of the one-sample Wilcoxon signed rank test for each decoder’s AUC against chance; * p < 0.05; ** p < 0.01; *** p < 0.001.

### Perception of decoder performance improves with tolerance for time discrepancies

To evaluate how participants’ perceived decoder performance would change according to tolerance for time discrepancies between their movement intention and stimulation onset, we analyzed decoder accuracy as a function of time window tolerance (**Figure 8a**). Predicted movement onsets were classified as true or false positives depending on whether they fell within or outside the tolerance window (**Figure 8b**). We observed that true positives and true negatives increased with increasing tolerance window, while false negatives and false positives decreased (**Figure 8c).**

**Figure 8:**
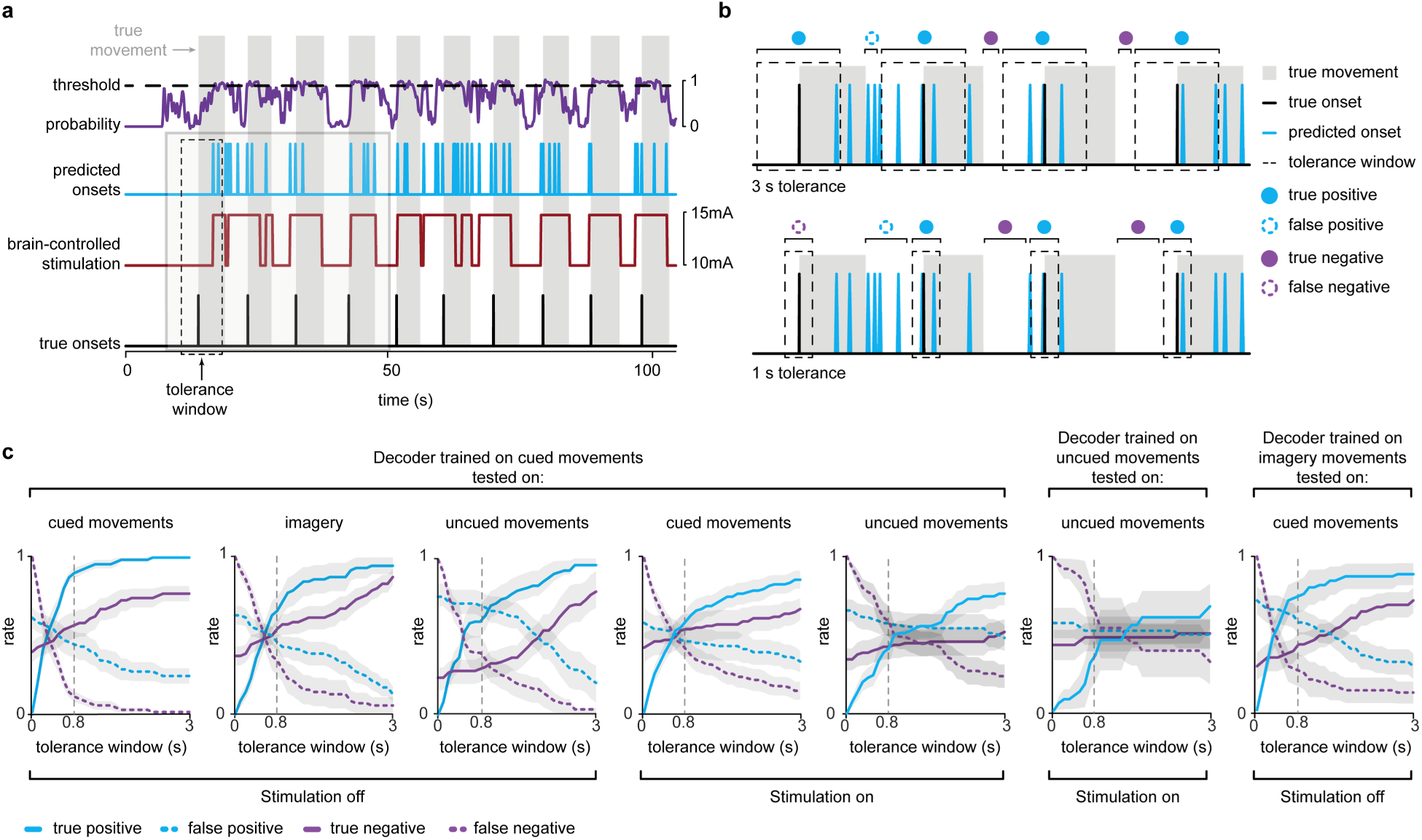
Decoder accuracy changes as a function of tolerance for discrepancies in time. **a.** Illustration of onset detection accuracy calculation. Onset detection accuracy was calculated as a function of tolerance time window length around true onset. Predicted onsets were defined as a positive crossing of the probability above threshold. **b.** Illustration of varying the tolerance window around true onset to calculate the onset detection accuracy. Tolerance window was varied from 0 to 3 seconds. True positives were considered any predicted onsets within the window, and true negatives were the absence of an onset prediction within rest. **c.** Onset detection accuracy plotted as a function of tolerance time window around true onset and averaged across participants for all conditions. The point of comparison across conditions (0.8 seconds) is denoted by a vertical dotted line.

To compare decoder performance across conditions, we used the tolerance window at which we observed diminishing returns on performance for the Phase I cued movement condition, which was at 0.8 seconds (**Figure 8c**). The mean of TPR and TNR with a 0.8-second tolerance window across conditions is reported in Supplementary Table 2. Importantly, we report the tolerance window as half of the total width, i.e., a tolerance window of 0.8 seconds implies onset ± 0.8 seconds for a full window length of 1.6 seconds. In Phase I, the mean of TPR and TNR at 0.8 seconds was 73% for cued movement (i.e., 73% of true onsets and rest periods are detected correctly with a 0.8-second tolerance window), 60% for imagery, and 44% for uncued movement. In phase I, the results of timing accuracy provide a conceptual understanding of how the decoder may perform with the addition of real-time tSCS.

In Phase II, the mean of TPR and TNR for cued and uncued movement were 56% and 41%, respectively (**Figure 8c**, **Supplementary Table 2**). This performance metric applied to real-time tested data implies that stimulation onset was within 0.8 seconds of true onset 56% of the time in cued movement and 41% of the time in uncued movement, demonstrating that the timing accuracy of movement onset prediction on uncued movement is generally lower than cued movement predictions. Combined with the AUC results, this implies that the decoder detects movement with above-chance accuracy when tested on both cued and uncued movements but performs with less accurate timing when tested on uncued movement.

The mean of TPR and TNR for a decoder trained and tested on uncued movement was 41% (**Figure 8c**, **Supplementary Table 2**), suggesting that the timing accuracy of stimulation onset does not tend to improve when training on uncued data. The mean of TPR and TNR for a decoder trained on imagery and tested on cued movement was 58% (**Figure 8c**). Therefore, the generalization between cued movement and imagery in Phase I has comparable timing accuracy to the generalization accuracy between imagery and cued movement in Phase III.

### Anthropomorphic cue is not essential for successful decoder performance

As a control and to determine whether action observation that activates motor regions related to the observed movement (26) was necessary to allow the accurate prediction of movement intention, we compared decoder performance when trained on anthropomorphic (knee cue) and non-anthropomorphic (bar cue) cues. Decoder performance for two participants during cued and uncued movement is shown in **Supplementary Figure 2**. Decoder performance tended to be similar when it was trained on the knee or bar cues, suggesting that an anthropomorphic cue was not essential for the ability of the decoder to accurately predict movement intention for the cued or uncued movement conditions.

## Discussion

### Summary

In this study, we validated a noninvasive BSI in unimpaired participants, with the aim of applying it as a rehabilitation strategy for motor recovery of lower limb function in individuals with SCI. Our findings demonstrate that the decoder can accurately predict the onset of lower limb movement through EEG signals better than chance and with consistent performance to similar decoders in the literature (21,36). This capability allows for the delivery of tSCS synchronized with movement intention in real time. Importantly, we verified that the decoder’s predictions were based on cortical activity rather than movement or cue-related artifacts, as evidenced by above-chance performance in imagined and uncued movement conditions. Furthermore, we demonstrated the decoder’s effectiveness in real-time control of tSCS, even in the presence of stimulation artifacts. These results suggest that this BSI system could be effectively integrated into rehabilitation strategies aimed at enhancing neuroplasticity and improving motor recovery in people with SCI.

### Generalization to imagery trials suggests similarities in neural strategies between tasks

Scalp EEG recordings have been reported as being highly susceptible to motion-related artifacts that can distort neural data (13,37,38). Our results show that the neural decoder trained on real, cued movements accurately generalized to the imagery condition (**Figure 3b, h, k**), and *vice versa (***Figure 7**). These observations suggest that detected event-related desynchronization in the µ (8-12 Hz), low β (16-20 Hz), and high β (24-28 Hz) frequency bands was related to movement intention rather than movement artifacts. Moreover, our results suggest that individuals employ a similar neural strategy when they imagine extending their knee and when they extend their knee, as demonstrated by consistent desynchronization topography maps across conditions (**Figure 4**). These results are in agreement with previous studies showing event-related desynchronization similarities between imagined and preparatory movement (15,16) and that imagery trials can be successfully used as training sets in BCIs that are used to control exoskeletons or paralyzed limbs (30–35). Furthermore, our results expand the potential application of our BSI to participants with no residual movement, given that sensorimotor event-related desynchronization is not substantially reduced due to cortical reorganization post-injury (39). In these cases, a training set consisting of imagery could be used in the delivery of tSCS timed with cued movement, enabling a participant with no residual movement to perform rehabilitation tasks.

### Cued and uncued movements may employ different neural strategies

The lower decoder performance in generalization from cued to uncued movements (**Figure 3c, i, l**) could be explained by widespread differences in the desynchronization patterns between these tasks (**Figure 4a, c, d**). Neural population activity in the motor cortex during reaching tasks has been shown to be consistent across direction, curvature, and distance, but varies with temporal dynamics such as velocity and reaction time (40,41). Additionally, neural firing in the dorsal premotor area (PMd) and supplementary motor area (SMA) has been shown to increase during voluntary self-paced movements (42,43), while putaminal activation occurs with a shorter latency during self-paced movement compared to cued movements (44). Together with our results, these observations suggest that cued and uncued movements may employ different neural strategies, creating important implications for BSIs that are aimed to be used during naturalistic, uncued movements.

### Generalization to naturalistic, uncued movements requires further improvements

Enabling the BSI to work under naturalistic, uncued movements would allow individuals with SCI to practice tSCS-assisted leg movements at their own pace and without the need to rely on an external cue. We investigated whether training our decoder on uncued movements could increase performance during the same task. While the average AUC increased (**Figure 6b**), the effect was not significant. Although this could be due to the low number of participants, our results indicate that the new decoder did not capture the strategy in uncued movement significantly better than a decoder trained on cued movements. Similarly, confusion matrices averaged across participants revealed poor performance of the stimulation paradigm (**Figure 6c**), suggesting that online predictions were inconsistent and empirically tuned probability thresholds could not account for unpredictable fluctuations in online probability.

The low performance across decoders trained on uncued movement could be due to several factors. Although we instructed participants to try to be as consistent with their movements as possible, we did not enforce this strategy. Therefore, the uncued task may contain higher variability in movement patterns compared to the cued condition, which negatively impacted decoder performance. Our subject-specific analysis revealed that desynchronization patterns were consistent across cued and uncued movements for some participants but not for others (**Figure 4a, Supplementary Table 3**). Although we did not analyze these populations separately due to the low number of participants, participants with consistent activation patterns may have higher generalization performance than those with inconsistent patterns. Desired neural patterns have been used as a screening tool for EEG-based BCI studies (30,31). However, we included all participants without *a priori* screening phase. Nevertheless, individuals can learn to generate the desired neural patterns via long-term BCI biofeedback training (12). Therefore, enabling the non-invasive BSI to work under naturalistic, uncued movements remains an important area of research for future work.

### Neural decoder limitations and potential improvements

An LDA algorithm was chosen in this study for its low variance, fast training/testing time, and interpretability of feature space (45). The assumptions of linearity, shared class covariance, and normality were not explicitly tested in this study. However, LDA classifiers have been shown to be robust to such violations in similar applications (21,36,45). Nonlinear models and neural networks could be explored to optimize performance (32,45), but the increased training time and training set size due to the complexity of these algorithms would require a significant increase in performance to justify their use (9). Moreover, as this technology progresses toward clinical applications, the recalibration time between sessions will become a significant factor. Determining the optimal recalibration intervals will be crucial to ensure consistent performance and adaptability of the system for individual patients over multiple sessions.

An analysis of the prediction performance at every timepoint at a particular probability threshold, as seen in the confusion matrices (**Figure 3j, k, l; Figure 5g, h; Figure 6c**, **Figure 7c**), showed a bias toward TPR over TNR, especially in Phase I. The probability threshold was empirically tuned by incorporating participants’ feedback in real time during Phase II. These results show that although we did not intend to have a bias one way or another, this empirical tuning strategy resulted in a bias toward TPR, suggesting that the probability thresholds were likely too low. Therefore, we reported an average of TPR and TNR as a balanced metric of decoder accuracy. In future developments, a mathematical optimization of the threshold on the probability could be performed when initially training, validating, and testing the decoder before it is applied in real-time tSCS control so that the true positives and true negatives would be balanced.

In applying BCIs in rehabilitation, an essential consideration is the delay between the evoked neural state and the biofeedback (46). In our BSI system, the primary factor contributing to the delay is the online low pass filter at 2Hz, which was added to ensure smooth decoder performance and allow enough time for real-time data processing. However, the delay in our system may need to be reduced to allow for Hebbian learning, in which the connections between simultaneously activated neurons are strengthened, allowing for neuroplasticity and synaptic reorganization in people with SCI (3,10,12). Applying tSCS precisely timed with the neural state evoked during movement could better improve motor outcomes as this system is applied as a rehabilitation strategy. Therefore, careful attention should be given to maintaining a balance between fast stimulation timing and decoder performance (**Figure 8**).

### Considerations for clinical translation involve further study of rehabilitative effects

While we customized the neural decoder according to each participant’s EEG data, an avenue for future research could be to test a generalized decoder trained on data from all participants. This approach could reveal whether a universal decoder can perform comparably to personalized ones, potentially simplifying the deployment process in clinical settings. However, one critical aspect to consider is that cortical representation of body areas may reduce in size or migrate after long-term loss of motor function (47–49). Sensorimotor regions containing event-related desynchronization in people with SCI may be different than those reported here, and unique for each participant (39,50–52).

BCI systems are thought to promote cortical plasticity, but the optimal method to induce neuroplasticity leading to motor recovery remains poorly understood. Previous applications of BCI systems in rehabilitation paired a desired neural state with FES feedback, training participants to recreate a predetermined neural state to receive FES supporting their movements, and thus inducing functional recovery (12). While shown to be effective in the rehabilitation of upper limb function in people with stroke, inducing a desired neural state rests on the assumption that neurological recovery involves restoring a state that resembles unimpaired controls. In our application, we train a decoder to identify consistencies in the existing neural state during movement and pair this neural state with tSCS feedback. While it is unclear which method would be better at promoting recovery, the question remains of how well a user can learn to control a decoder. Nevertheless, co-adaptation of the user and BCI system will likely be essential to achieve natural and improved control (9,53).

## Conclusion

In this work, we demonstrate the feasibility of a noninvasive BSI in unimpaired participants. The accurate decoding of EEG signals can be used to synchronize tSCS with movement intention and shows promise for improving neurorehabilitation techniques. Continued development and optimization of this system could further enhance its efficacy and accessibility. Integrating this BSI system into rehabilitation protocols could enhance neuroplasticity and motor recovery, while also advancing our understanding of BCI-based rehabilitation.

## Acknowledgements

This work was supported primarily by the McDonnell Center for Systems Neuroscience at Washington University in St. Louis. C.A., L.L., R.K., R.H., and I.S. received partial support from the National Institutes of Health NICHD Award Number K12-HD073945 and NINDS Award Numbers K01-NS127936. E.C.L., P.B. received partial support from U24-NS109103, NIBIB Award Numbers R01-EB026439 and P41-EB018783, and internal funding from the Department of Biomedical Engineering and the Department of Neurosurgery.

The authors thank Noah Bryson for the software development related to the data collection for recruitment curves, Phillip Demarest, Jürgen Mellinger, Nicholas Luzcak, James Swift, and Katrin Mayr for BCI2000 software adaptation for this study, and Dr. Simon Borgognon for statistical analysis code development.

## Author contributions

C.A., and L.L, data analysis.

C.A., L.L., M.L. and Z.S., conducted experiments. L.L, M.L, and R.K., software development.

E.C.L., P.B. and I.S., conceptualization and supervision.

C.A., L.L, M.L., R.K., R.H., Z.S., E.C.L., P.B., and I.S., data interpretation.

C.A., L.L., and I.S., manuscript writing, review, and editing.

## Competing interests

E.C.L. holds various patents in relation to the present work and is a founder and shareholder of Neurolutions, Inc., a company developing EEG technologies for stroke. All other authors declare no competing interests.

## Data availability

Data from this study will be made available upon reasonable request to the corresponding author.

## Code availability

All software used to produce the figures in this manuscript will be available upon reasonable request to the corresponding author.

## Supplementary Data

**Supplementary Figure 1.**
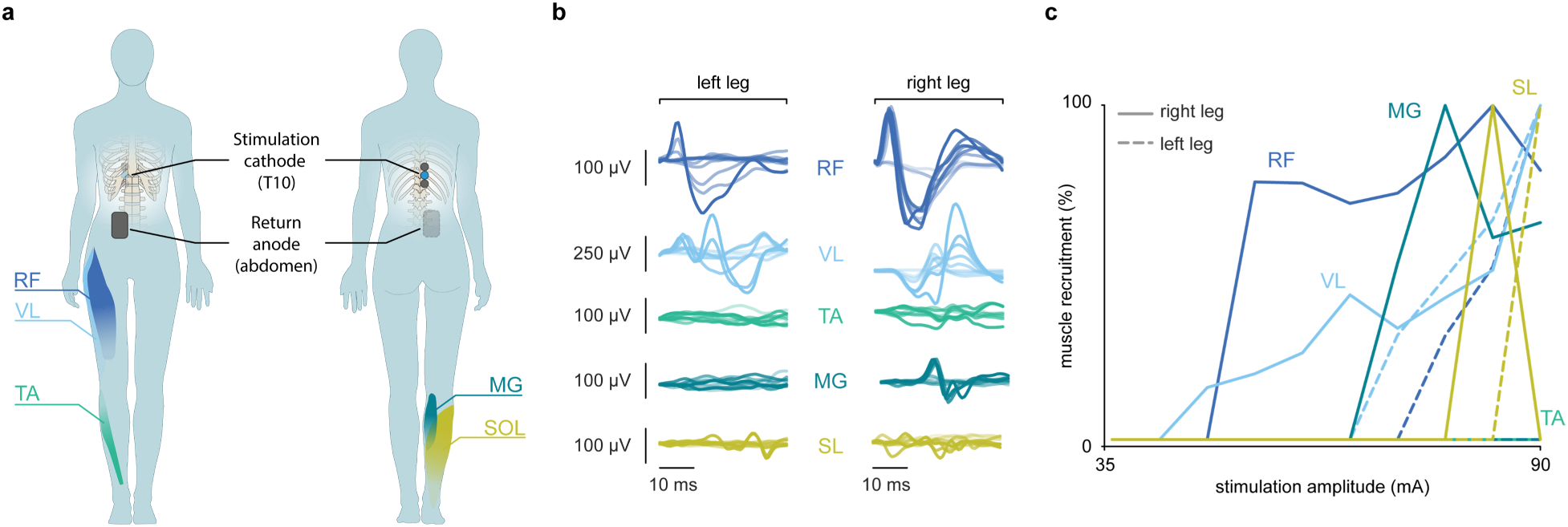
Stimulation protocol to selectively target right hip muscles. **a.** Configuration of tSCS. Placement of the stimulation cathode at spinal segment T10 and return anode to the right of the naval to selectively target right hip muscles. **b.** Muscle responses. Muscle responses from major leg muscles for pulses of increasing amplitudes. **c.** Recruitment curves. Muscle response peak-to-peak amplitude as a function of stimulation amplitude. Abbreviations: rectus femoris (RF); vastus lateralis (VL); tibialis anterior (TA); medial gastrocnemius (MG); soleus (SL).

**Supplementary Figure 2.**
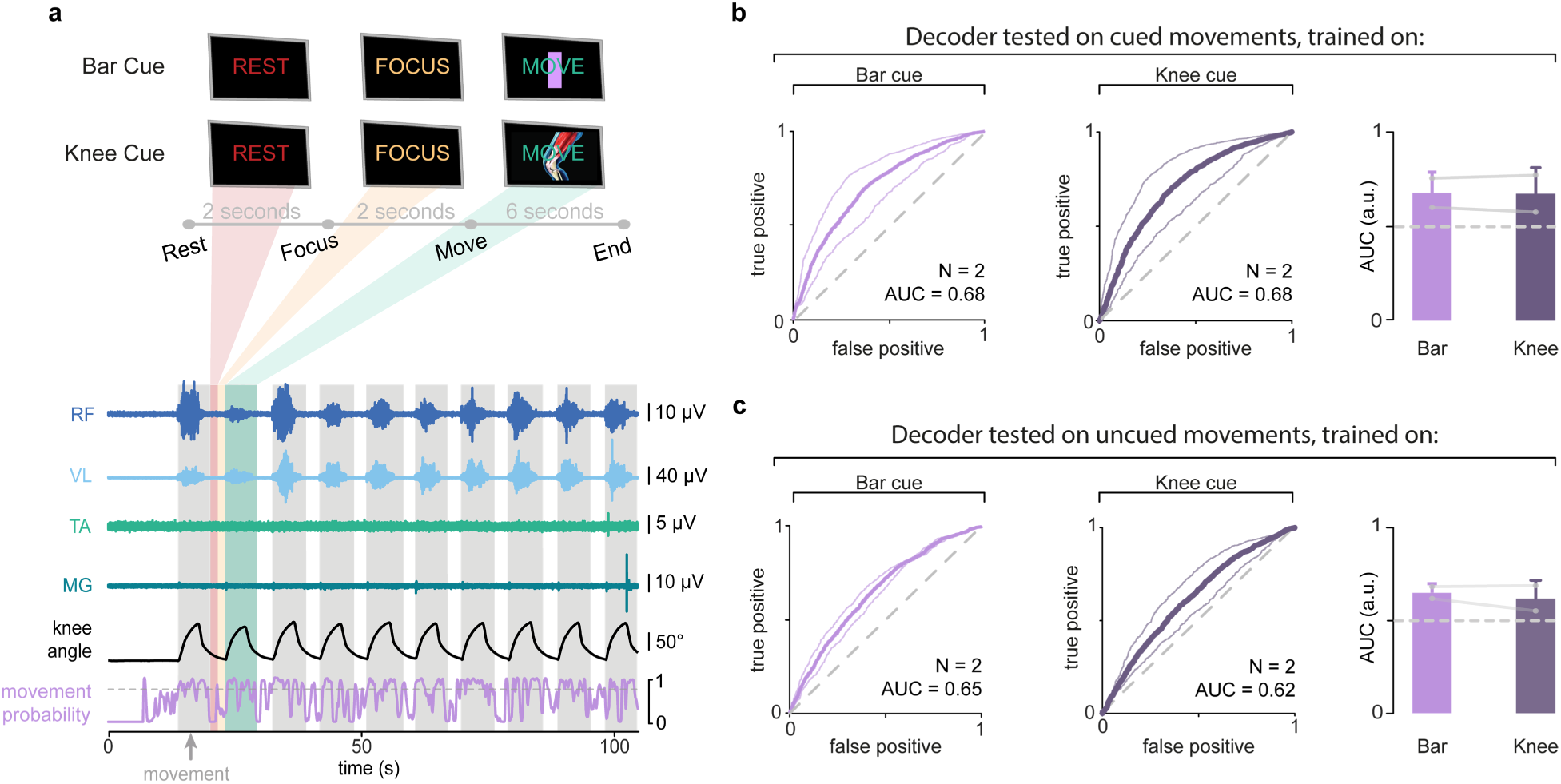
Anthropomorphic cue is not necessary for successful decoder performance. **a.** Illustration of cue types within the training sets. **b.** ROC curve for all participants (thin line) and averaged across participants (thick line) when the decoder was trained on the bar and knee cues and tested on cued movements. **c.** ROC curve for all participants (thin line) and averaged across participants (thick line) when the decoder was trained on bar and knee cues and tested on uncued movement.

**Supplementary Table 1.** Participant demographics. In column 3, I. indicates the uncued movement-based decoder experiment, and II. indicates the anthropomorphic vs. bar cue experiment. *Datasets excluded from analysis due to outliers in channel noise.

| Participant | Age | Sex | Phase I | Phase II | Phase III |
| --- | --- | --- | --- | --- | --- |
| 001 | 35 | M | Yes | Yes | No |
| 005 | 29 | M | Yes | No | No |
| 006 | 25 | M | Yes | Yes | II. |
| 007 | 33 | F | Yes | No | No |
| 008 | 26 | F | Yes | Yes | I. |
| 009 | 22 | F | Yes | No | No |
| 010 | 27 | M | Yes* | No | No |
| 011 | 19 | M | Yes | No | No |
| 013 | 23 | F | No | Yes | I., II. |
| 014 | 29 | M | No | Yes | No |
| 015 | 27 | M | No | Yes | No |
| 016 | 23 | M | No | Yes | No |
| 017 | 26 | F | No | Yes* | No |
| 018 | 23 | F | No | Yes | No |
| 019 | 24 | M | No | Yes | No |
| 020 | 28 | M | No | No | I. |
| 021 | 23 | F | No | No | I. |

**Supplementary Table 2.** Onset detection performance comparison at a tolerance window of 0.8 seconds around true onset. This tolerance window was the inflection point of the derivative of the onset detection TPR curve for Phase I cued movement and applied across conditions for comparison of results. Values are presented as percentages.

| Onset Detection Performance |  |  | Average |
| --- | --- | --- | --- |
| Phase I |  |  |  |
| Cued | TPR | 89.1 | 72.5 |
|  | TNR | 55.9 |  |
| Imagery | TPR | 65.5 | 59.8 |
|  | TNR | 54.0 |  |
| Free | TPR | 60.5 | 44.4 |
|  | TNR | 28.3 |  |
| Phase II |  |  |  |
| Cued | TPR | 58.5 | 55.9 |
|  | TNR | 53.2 |  |
| Free | TPR | 40.1 | 40.8 |
|  | TNR | 41.5 |  |
| Phase III |  |  |  |
| Free (trained/tested) | TPR | 35.1 | 41.3 |
|  | TNR | 47.5 |  |
| Cued (trained on Imagery) | TPR | 74.2 | 57.6 |
|  | TNR | 40.9 |  |

**Supplementary Table 3.**
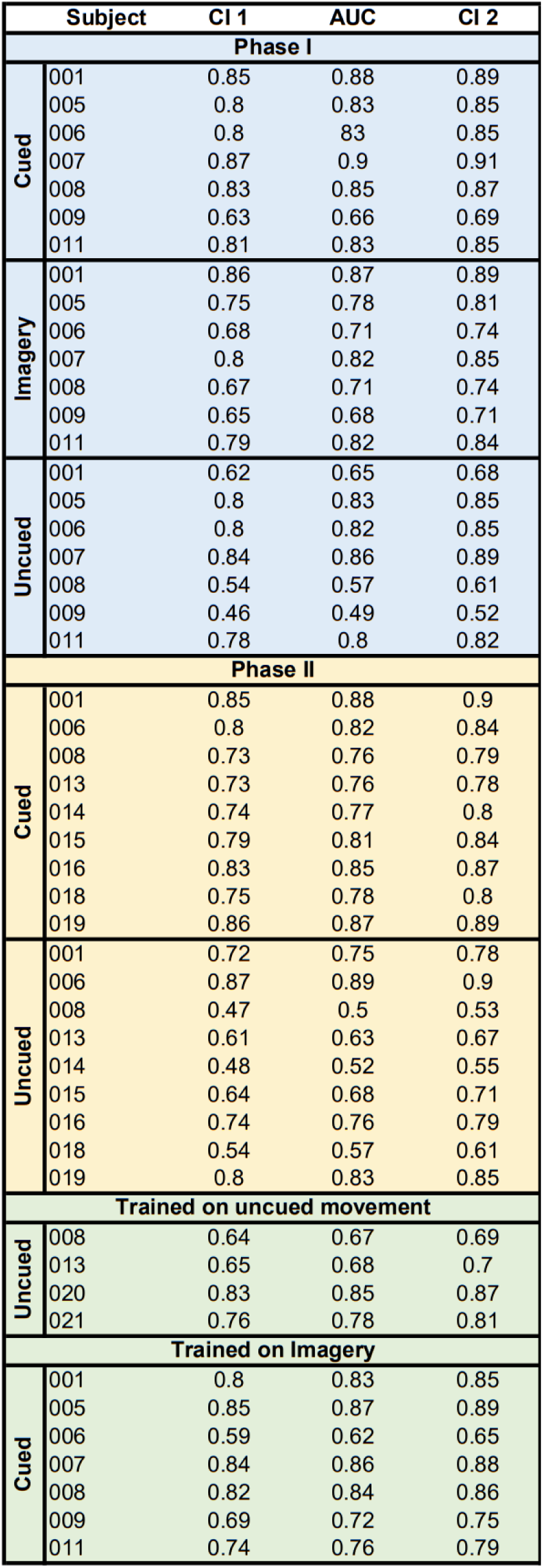
Decoder AUC and confidence intervals. AUCs for each participant’s decoder were calculated from the original test set. The 95% confidence intervals were constructed from bootstrapping.

